# Suppression of ferroptosis by vitamin A or antioxidants is essential for neuronal development

**DOI:** 10.1101/2023.04.05.535746

**Authors:** Juliane Tschuck, Vidya Padmanabhan Nair, Ana Galhoz, Gabriele Ciceri, Ina Rothenaigner, Jason Tchieu, Hin-Man Tai, Brent R. Stockwell, Lorenz Studer, Michael P. Menden, Michelle Vincendeau, Kamyar Hadian

## Abstract

Development of functional neurons is a complex orchestration of several signaling pathways controlling cell proliferation, differentiation, and homeostasis^1^. However, details about the involved factors are not fully understood. The balance of antioxidants and vitamins is important for neuronal survival, synaptic plasticity, and early neuronal development; thus, we hypothesized that ferroptosis—a lipid peroxidation dependent cell death modality that is inhibited by antioxidanats^2,3^—needs to be suppressed to gain neurons. Our study shows that removal of antioxidants diminishes neuronal development and laminar organization of cortical organoids. Intriguingly, impaired neuronal development in conditions lacking antioxidants can be fully restored when ferroptosis is specifically inhibited by ferrostatin-1, or neuronal differentiation occurs in the presence of sufficient amounts of vitamin A. Mechanistically, vitamin A activates the heterodimeric nuclear receptor complex Retinoic Acid Receptor (RAR)/Retinoid X Receptor (RXR)^4^, which upregulates expression of the ferroptosis regulators GPX4, FSP1, GCH1, and ACSL3, amongst others. Therefore, our study reveals that above a certain threshold, vitamin A increases expression of essential cellular gatekeepers of lipid peroxidation. This study uncovers a critical process during early neuronal development, where suppression of ferroptosis by radical-trapping antioxidants or vitamin A is required to obtain maturing neurons and proper laminar organization in cortical organoids.

## Introduction

During brain development, neurons arise in a tightly controlled process regulating an important balance between proliferation and differentiation events^1^. To ensure that cells emerge, migrate and mature at the right time and space, a network of genetic and molecular factors is required^5^. However, the exact factors and their precise interplay are still not fully understood. Regulated cell death (RCD) eliminates half of the neurons initially generated during mammalian brain development and helps shaping the organization of cortical circuits by regulating their cellular composition^6,7^. Apoptosis, a well-studied RCD modality, has so far been described as the major form of cell death during the developing nervous system^6,7^.

Ferroptosis is another RCD, which occurs through iron-dependent lipid peroxidation^3,8^. It is controlled by a number of cellular gatekeepers including the system-xc^-^/glutathione peroxidase 4 (GPX4)/glutathione axis^3^, the ferroptosis suppressor protein 1 (FSP1)/ubiquinol/vitamin k axis^9–11^, the GTP cyclohydrolase 1 (GCH1)/tetrahydrobiopterin (BH4)/dihydrofolate reductase (DHFR) axis^12,13^, and the nuclear receptor Farnesoid X receptor (FXR)^14,15^, amongst others. Ferroptosis can be the cause for several degenerative diseases of the adult brain, heart, lung and kidney^3^. However, our understanding about the functions of ferroptosis during brain development remains rudimentary. In this study, we illuminate the detrimental role of increased ferroptosis during neurogenesis and the necessity to inhibit ferroptosis by radical-trapping antioxidants or vitamin A to obtain cortical neurons.

## Results

### Vitamin A restores neuronal differentiation in the absence of antioxidant protection

Most neuronal differentiation protocols require a series of antioxidants (AO) (Supplementary Table 1) including vitamin E and glutathione (GSH), two well-known ferroptosis inhibitors^2^, for proper neuronal development. Thus, to understand whether exclusion of these antioxidants would affect neurogenesis and whether vitamin A could compensate for the antioxidant deficiency, we differentiated human embryonic stem cells (H9; WA09) into cortical neurons using an established differentiation protocol^16–18^. It is reported that the concentration of all-trans retinoic acid (ATRA) (active vitamin A metabolite) is important for its distinct functions^19,20^. Hence, we used three conditions without or with increasing vitamin A concentrations between day-10 to day-20 of differentiation (**Fig. 1a**): (i) standard media with antioxidant-containing B27 without vitamin A (+AO; Supplementary Table 1), (ii) media with antioxidant-deficient B27 containing vitamin A (-AO+vA^lo^; Supplementary Table 1), or (iii) media with antioxidant-deficient B27 containing vitamin A, which was further supplemented with ATRA (-AO+vA^hi^). We performed total RNA sequencing of the +AO and -AO+vA^lo^ samples and showed that vitamin A supplementation led to overexpression of Retinoic Acid Receptor B (RARB) and beta-carotene oxygenase 2 (BCO2) at day-20 (immature neurons) in -AO+vA^lo^ treated cells compared to +AO treated cells (**Extended Data Fig. 1a,b**). RARB is a nuclear receptor employing vitamin A (ATRA) as a ligand for transcriptional activity^4^, and BCO2 is an enzyme involved in retinoic acid biosynthesis^21^. Increased expression of RARB and BCO2 at day-20 in -AO+vA^lo^ treated cells could be validated using qRT-PCR (**Extended Data Fig. 1c**). We also checked levels of RARB upon low and high exposure to vitamin A and saw a dose-dependent increase in RARB expression (**Fig. 1b**). Interestingly, lack of antioxidants had the consequence that immature neurons at day-20 of differentiation showed higher lipid peroxidation as detected by the C11-BODIPY lipid peroxidation sensor. However, higher levels of vitamin A could fully abrogate lipid peroxidation (**Fig. 1c**). We further differentiated the three conditions into day-40 cortical neurons and demonstrate that -AO+vA^lo^ treated conditions showed significantly fewer MAP2-positive neurons (**Fig. 1d**). MAP2a/b isoforms are neuron*-*specific cytoskeletal proteins and a suitable marker for neuronal cells to investigate morphological (cytoskeletal) changes^18^. However, higher levels of vitamin A (-AO+vA^hi^) restored the number of MAP2-positive cortical neurons (**Fig. 1d**). Interestingly, those neurons that were generated under -AO+vA^lo^ conditions showed similar levels of axon length, branching and extremities (**Extended Data Fig. 2a**), arguing for cell death under this condition due to lack of antioxidants rather than impaired neuronal differentiation.

**Fig. 1:**
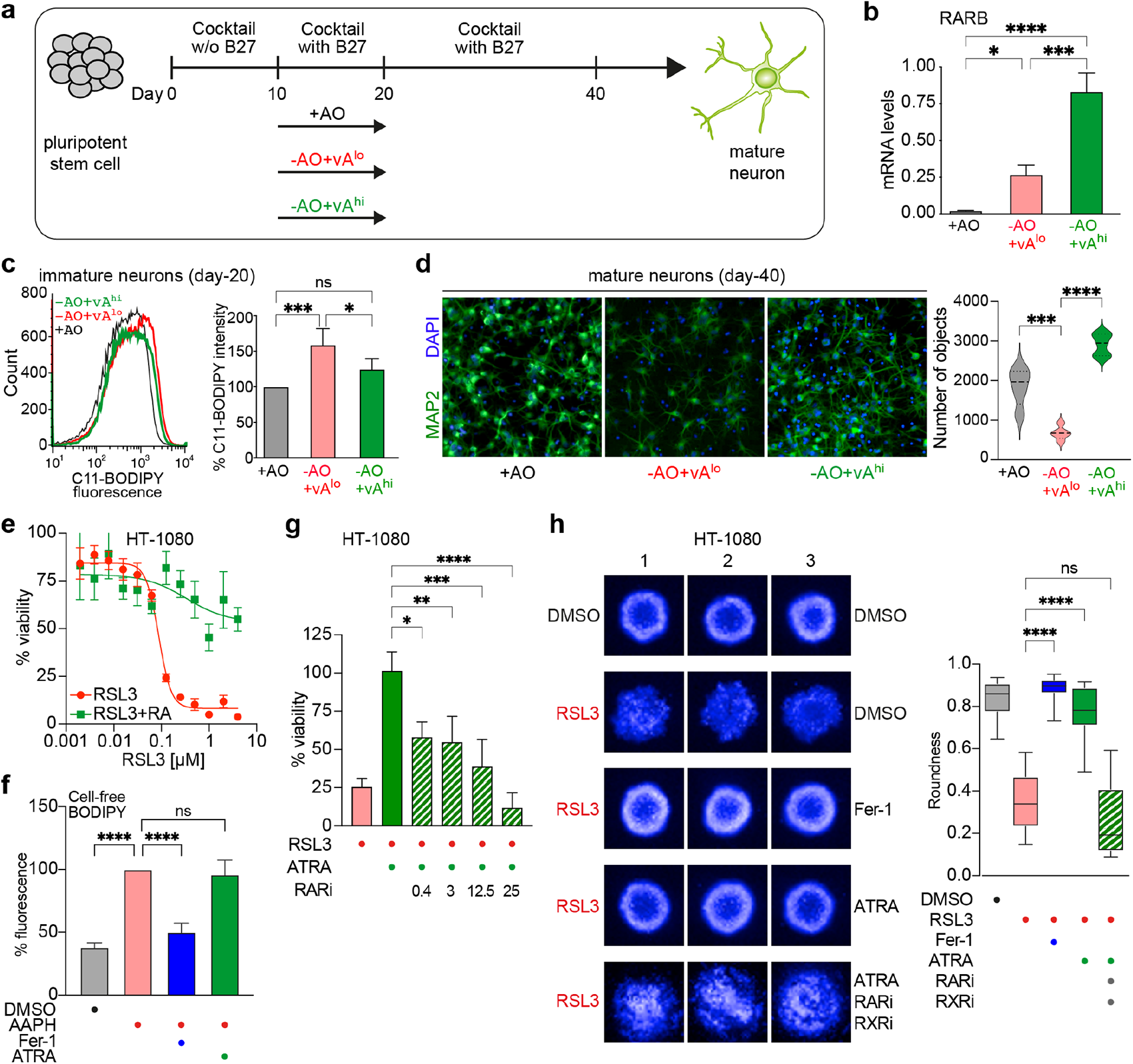
Vitamin A restores neuronal differentiation by reducing lipid peroxidation. **a**, Scheme of neuronal differentiation using three conditions (i) standard media with antioxidant-containing B27 (+AO), (ii) media with antioxidant-deficient B27 containing vitamin A (-AO+vA^lo^), or (iii) media with antioxidant-deficient B27 containing vitamin A, which was further supplemented with retinoic acid (RA) (-AO+vA^hi^). **b**, mRNA levels of RARB at day 20 of cortical neuronal differentiation using quantitative RT-PCR (n=3). Data plotted are mean ± SD. **** p≤ 0.0001, *** p≤ 0.001, * p≤ 0.05 (One-way-ANOVA). **c**, C11-BODIPY staining of day-20 immature cortical neurons supplemented with vitamin A using flow cytometry (n=3). (Left) flow cytometry histograms; (right) median intensity—data plotted are mean ± SD; *** p≤ 0.001, * p≤ 0.05 (One-way-ANOVA). **d**, (Left), MAP2 immunofluorescence staining of cortical neurons at day 40. (Right), high-content image analyses (n=3). Data plotted are mean ± SD. **** p≤ 0.0001, *** p≤ 0.001 (One-way-ANOVA). **e**, CellTiter-Glo assay of HT-1080 cells co-treated with the ferroptosis inducer RSL3 and vitamin A (ATRA) (n=3). **f**, C11-BODIPY cell-free assay of the radical initiator 2,2’-Azobis(2-amidinopropane) dihydrochloride (AAPH) co-treated with the radical-trapping antioxidant Fer-1 (Fer-1) or vitamin A (ATRA) (n=3). Data plotted are mean ± SD. **** p≤ 0.0001 (One-way-ANOVA). **g**, CellTiter-Glo assay of HT-1080 cells after co-treatment with RSL3 and vitamin A (ATRA) as well as different concentrations of the small molecule inhibitor of Retinoic Acid Receptor (RARi) (n=3). Data plotted are mean ± SD. **** p≤ 0.0001, *** p≤ 0.001, ** p≤ 0.01, * p≤ 0.05 (One-way-ANOVA). **h**, (Left), HT-1080 derived spheroids co-treated with the ferroptosis inducer RSL3 and ferroptosis inhibitor Fer-1, vitamin A (ATRA), or in combination with small molecule inhibitors of Retinoic Acid Receptor (RARi) and Retinoid X receptor (RXRi). (n=13; remaining replicates of spheroids in Extended data Fig. 3) (Right), quantification of spheroid roundness. **** p≤ 0.0001 (One-way-ANOVA); n=3 refers to biologically independent experiments.

### Vitamin A inhibits ferroptosis by reducing lipid peroxidation

Next, we aimed at understanding the mechanism by which vitamin A counteracts ferroptotic cell death, and whether the effect is specific to neurons. Hence, we used the fibrosarcoma cell line HT-1080 that is well-established to study ferroptosis^8,10,12^. In an ATP-based viability assay, cells were treated with the ferroptosis inducer (FIN) RSL3 and co-treated with vitamin A (ATRA). ATRA significantly rescued cells from ferroptotic cell death (**Fig. 1e** and **Extended Data Fig. 2b,d**). A similar effect was achieved when ferroptosis was induced by the system xc^-^ inhibitor IKE (**Extended Data Fig. 2c**). Moreover, the anti-ferroptotic impact of ATRA was similar to ferrostatin-1 (Fer-1), a gold standard ferroptosis inhibitor^8^ (**Extended Data Fig. 2b,c**).

To elucidate whether this ferroptosis-inhibiting effect of vitamin A can be ascribed to a possible direct antioxidative property, we performed several cell-free assays: first, oxidation of BODIPY 581/591 C11 was induced by 2,2’-Azobis(2-amidinopropane) dihydrochloride, a free radical initiator. Addition of the radical-trapping antioxidant Fer-1 could stop the oxidation reaction, whereas addition of ATRA had no effect (**Fig. 1f**). In a second antioxidation assay, we initiated a radical reaction using 2,2-diphenyl-1-picrylhydrazyl (DPPH), and again Fer-1 acted as a radical scavenger, but ATRA showed no activity (**Extended Data Fig. 2e**). These data, in the absence of the cellular environment, suggest that the ferroptosis-inhibiting effect of vitamin A is not caused by any direct antioxidative properties. All-trans retinoic acid (ATRA) activates the RAR nuclear receptors to initiate gene transcription, raising the possibility that vitamin A inhibits ferroptosis via transcriptional control. To test this hypothesis, we co-treated HT-1080 cells with ATRA and small molecule inhibitors of Retinoic Acid Receptor (RARi, AGN193109) or Retinoid X receptor (RXRi, HX531), respectively, and observed that both inhibitors re-sensitized cells to RSL3-mediated ferroptosis in the presence of ATRA (**Fig. 1g**, Extended Data Fig. 2f).

We also cultured HT-1080 spheroids to validate these findings in a 3D model (**Fig. 1h** and **Extended Data Fig. 3**). We induced ferroptotic cell death using RSL3, which consequently prevented the formation of spheroids. Treatment with ATRA restored the round shape of the spheroids to the same extent as Fer-1. Interestingly, co-treatment of ATRA with RARi/RXRi re-sensitized spheroids to RSL3 destruction, leading to loss of roundness as a result of ferroptotic cell death. These results were quantified by performing a Hoechst 33342 stain with subsequent high-content-image analysis that calculated spheroid roundness (**Fig. 1h**).

Next, we investigated whether vitamin A, although without direct antioxidative function, would be able to reduce lipid peroxidation in a series of cell-based lipid peroxidation assays. First, we performed a TBARS assay that measures levels of malondialdehyde (MDA), a product of lipid peroxidation, and found that ATRA significantly reduced MDA levels similar to Fer-1 indicating a reduction of lipid peroxidation (**Fig. 2a**). Further, in a live cell staining with the fluorescent lipid peroxidation sensor C11-BODIPY 581/591 and subsequent flow cytometry, we observed that ATRA decreased RSL3-induced lipid peroxidation (**Fig. 2b**). We also performed C11-BODIPY 581/591 in microscopy imaging and found reduction of lipid peroxidation by ATRA (**Extended Data Fig. 4a**). Finally, by immunostaining and flow cytometry, another RSL3-induced product of lipid peroxidation in cell membranes, namely 4-hydroxynonenal (4-HNE), was found to be significantly reduced upon ATRA treatment (**Fig. 2c**). Hence, these studies demonstrate that vitamin A addition eliminates lipid peroxidation.

**Fig. 2:**
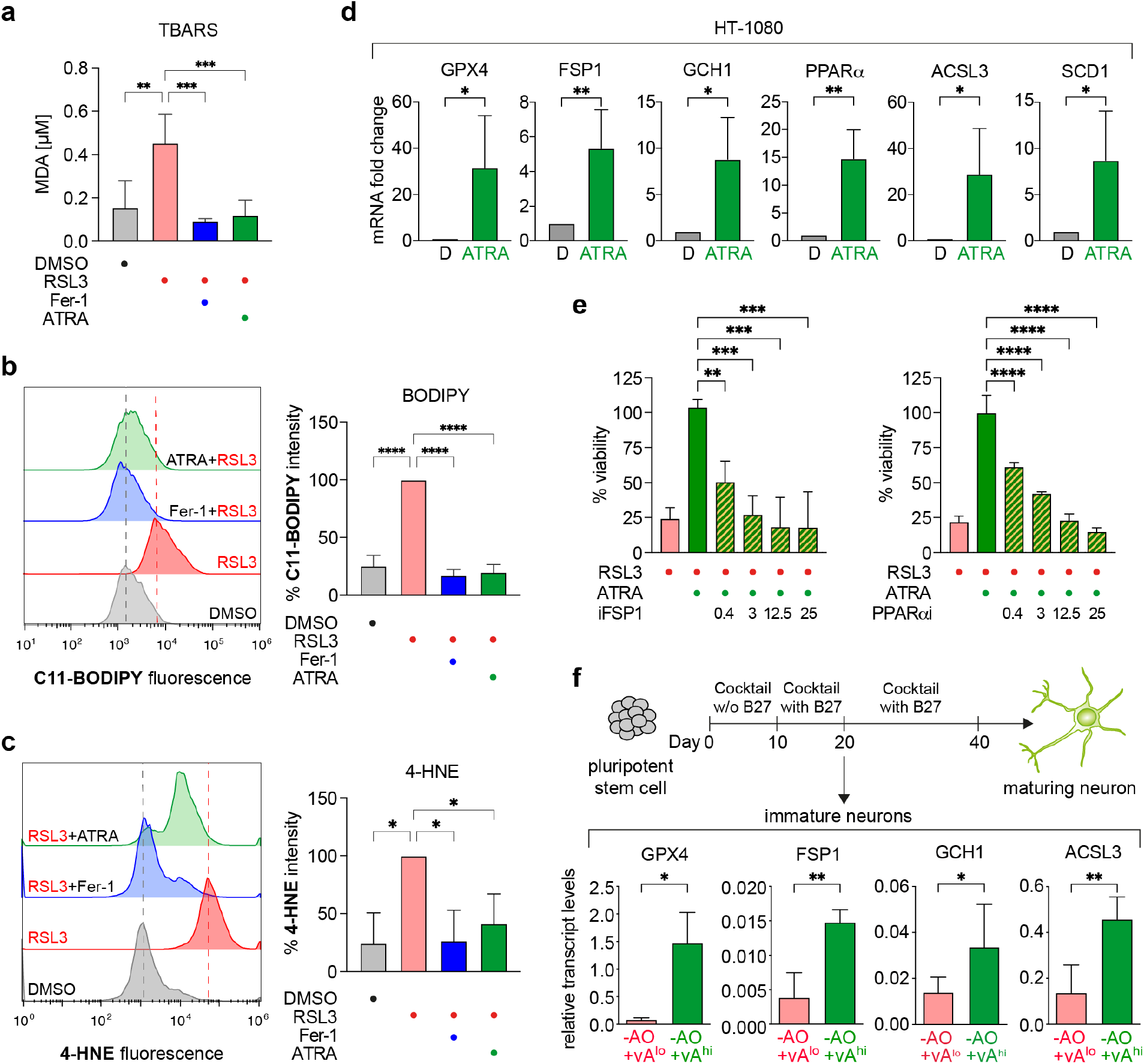
Vitamin A eliminates lipid peroxidation and upregulates major anti-ferroptotic regulators in HT-1080 as well as cortical neurons. **a**, TBARS assay of HT-1080 cells co-treated with the ferroptosis inducer (RSL3) and inhibitor (Fer-1) or vitamin A (ATRA) (n=3). Data plotted are mean ± SD. *** p≤ 0.001, ** p≤ 0.01 (One-way-ANOVA). **b**, Flow cytometry of C11-BODIPY staining: (left) histograms and (right) intensity of HT-1080 cells co-treated with the ferroptosis inducer (RSL3), and inhibitor (Fer-1) or vitamin A (ATRA) (n=3). Data plotted are % intensity of median fluorescence normalized to RSL3-treated cells ± SD. **** p≤ 0.0001 (One-way-ANOVA). **c**, Immunostaining of 4-Hydroxynonenal (4-HNE) and flow cytometry: (left) histograms and (right) intensity of HT-1080 cells co-treated with the ferroptosis inducer (RSL3), and inhibitor (Fer-1) or vitamin A (ATRA) (n=3). Data plotted are % intensity of median fluorescence normalized to RSL3-treated cells ± SD. * p≤ 0.05 (One-way-ANOVA). **d**, mRNA expression, using quantitative RT-PCR, of various anti-ferroptotic regulators in HT-1080 cells treated with DMSO or vitamin A (ATRA) (n=3). Data plotted are mean ± SD. ** p≤ 0.01, * p≤ 0.05 (unpaired t-test). **e**, CellTiter-Glo assay of co-treated HT-1080 cells with RSL3, vitamin A (ATRA) and inhibitors against FSP1 (iFSP1) or PPAR*α* (GW6471) (n=3). Data plotted are mean ± SD. **** p≤ 0.0001, *** p≤ 0.001, ** p≤ 0.01 (One-way-ANOVA). **f**, mRNA expression (qRT-PCR) of various anti-ferroptotic regulators in cortical neurons treated with low or high vitamin A (n=3). Data plotted are mean ± SD. ** p≤ 0.01, * p≤ 0.05 (unpaired t-test); n=3 refers to biologically independent experiments.

### Vitamin A upregulates major anti-ferroptotic regulators

Based on these findings that vitamin A activates RAR/RXR to reduce lipid peroxidation, we hypothesized that ATRA may alter expression of distinct ferroptosis-inhibiting regulators involved in suppression of lipid peroxidation. To verify this, quantitative RT-PCR experiments were performed in HT-1080 cells treated with ATRA (**Fig. 2d**). Several genes that are established as key ferroptotic gatekeepers were significantly upregulated upon ATRA treatment, *i.e.,* GPX4, FSP1, and GCH1—the three major pillars of anti-ferroptotic defense. Additionally, expression of anti-ferroptotic regulators involved in lipid biosynthesis and lipid metabolism were increased: Peroxisome Proliferator-Activated Receptor alpha (PPAR*α*), Acyl-CoA Synthetase Long-chain family member 3 (ACSL3), and Stearoyl-CoA desaturase (SCD1). We further co-treated HT-1080 cells with RSL3, ATRA and inhibitors against FSP1 (iFSP1) or PPAR*α* (GW6471) (**Fig. 2e**). Cotreatment with these inhibitors resulted in a loss of the anti-ferroptotic effect of ATRA to re-sensitize the cells towards ferroptosis; thus, proving these target genes are indeed downstream target of RAR/RXR activation.

We then evaluated whether this mechanism was also responsible for the anti-ferroptotic effect seen in immature neurons and performed quantitative RT-PCR of neurons at day 20 of differentiation, which were grown without antioxidants, and either supplemented with lower or higher levels of vitamin A (**Fig. 2f**). Only higher vitamin A levels upregulated GPX4, FSP1, GCH1, ACSL3, and SCD1 (**Fig. 2f, Extended Data Fig. 4b**), thereby explaining the protective effect of vitamin A during stem cell differentiation by reducing lipid peroxidation and ferroptosis (**Fig. 1d**). PPAR*α* was also upregulated to some extent upon higher vitamin A treatment, but was not significant (**Extended Data Fig. 4b**).

Thus, we report an unexpected function of vitamin A and its active component ATRA in inhibiting lipid peroxidation by enhancing the transcription of essential anti-ferroptotic defense modules, including the antioxidative enzymes GPX4, FSP1 and GCH1, as well as the lipid composition regulators ACSL3, SCD1, and PPAR*α*.

### Specific inhibition of ferroptosis promotes neuronal differentiation

After having established that vitamin A is a potent inhibitor of ferroptosis and sufficient to facilitate cortical neuronal differentiation in the absence of antioxidants, we aimed at further testing whether cells indeed undergo ferroptosis in the absence of antioxidants by using an established ferroptosis inhibitor, *i.e.,* ferrostatin-1 (Fer-1)^3,8^. Hence, we differentiated H9 cells into cortical neurons again using three differentiation conditions (**Fig. 3a**). Between day 10 to 20 of differentiation, we used (i) the standard B27-supplemented media containing antioxidants but not vitamin A (+AO), media with antioxidant-deficient B27 containing vitamin A (-AO+vA^lo^), or (iii) media with antioxidant-deficient B27 containing vitamin A, which was supplemented with ferrostatin-1 (-AO+vA^lo^+Fer-1) (**Fig. 3a**). At day 20 of cortical neuronal differentiation, we investigated the level of lipid peroxidation using the C11-BODIPY sensor (**Fig. 3b**). Cells that differentiated without antioxidants (-AO+vA^lo^) showed an increase of lipid peroxides compared to immature neurons differentiated under standard conditions (+AO) (**Fig. 3b**). Importantly, treatment of the cells with ferrostatin-1 (-AO+vA^lo^+Fer-1) reverted lipid peroxidation to control levels (**Fig. 3b**). Moreover, we differentiated the three conditions into maturing cortical neurons. Day-40 cortical neurons were analyzed for neuronal morphology using MAP2 immunofluorescence staining (**Fig. 3c**). Neurons differentiated without antioxidants and low levels of vitamin A (-AO+vA^lo^) showed a decrease in MAP2 expression as well as less neuronal segments when compared to neurons differentiated in the presence of antioxidants (+AO) (**Fig. 3c**). Addition of Fer-1 to the differentiation without antioxidants (-AO+vA^lo^+Fer-1) fully rescued the observed phenotype (**Fig. 3c**). Importantly, the number of healthy cells was reduced without antioxidants. As the ferroptosis-specific inhibitor Fer-1 fully restored MAP2 positive cells, these data suggest the occurrence of ferroptotic cell death during early events of differentiation in the absence of antioxidants rather than proliferative effects. Analysis of *MAP2* mRNA expression of day-40 cortical neurons confirmed reduced *MAP2* mRNA levels in -AO+vA^lo^ treated cells and a rescue of *MAP2* mRNA levels upon Fer-1 treatment (-AO+vA^lo^+Fer-1) (**Fig. 3d**).

**Fig. 3:**
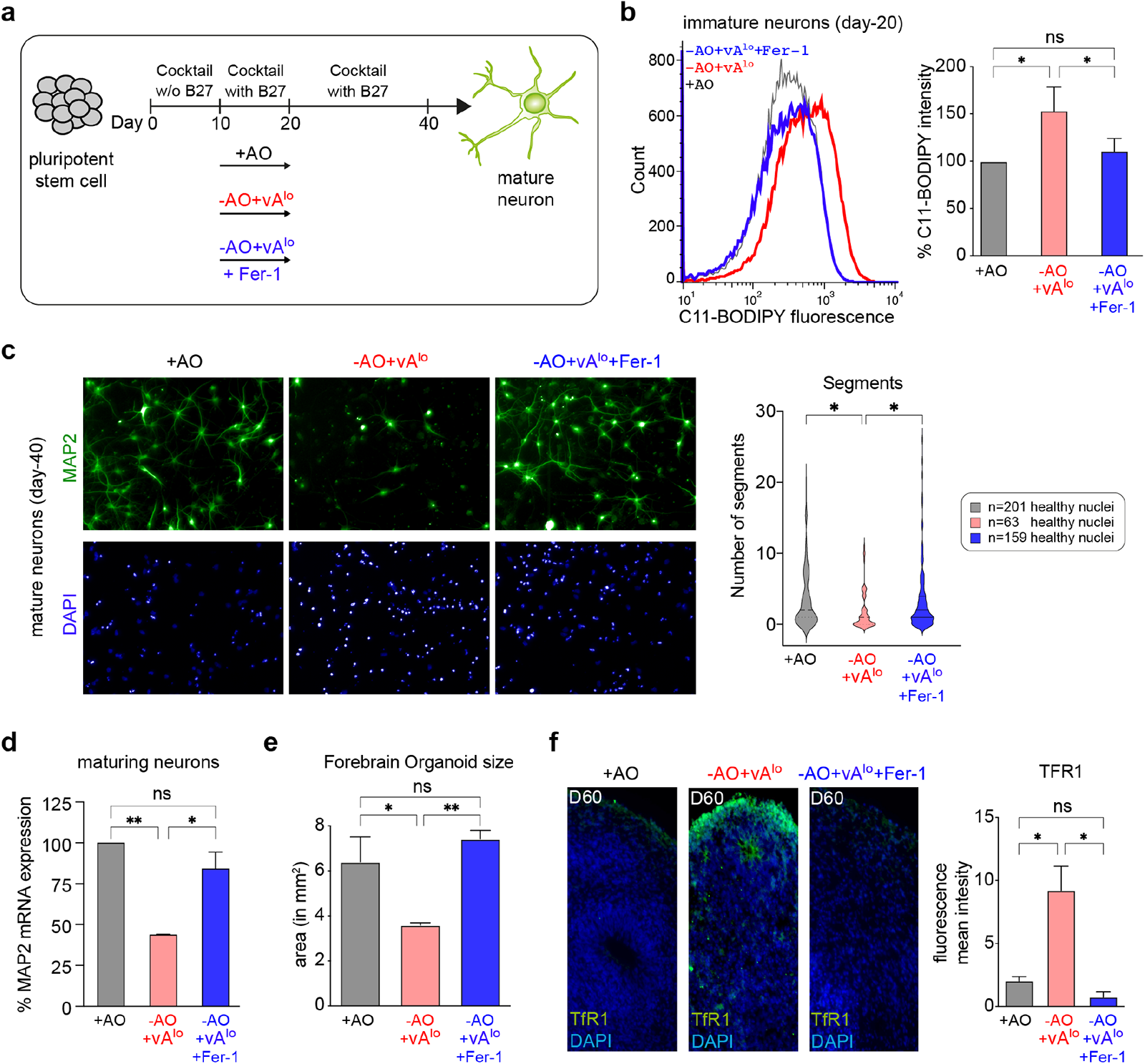
Inhibition of ferroptosis by ferrostatin-1, upon deprivation of antioxidants, facilitates neuronal differentiation. **a**, Scheme of cortical neuronal differentiation using three conditions (i) standard media with antioxidant-containing B27 (+AO), (ii) media with antioxidant-deficient B27 containing vitamin A (-AO+vA^lo^), or (iii) media with antioxidant-deficient B27 containing vitamin A, which was further supplemented with ferrostatin-1 (Fer-1) (-AO+vA^lo^+Fer-1). **b**, C11-BODIPY staining of day-20 immature cortical neurons generated with antioxidant (+AO) or without antioxidants (-AO+vA^lo^) or supplemented with ferrostatin-1 (-AO+vA^lo^+Fer1) using flow cytometry (n=3). (Left) flow cytometry histograms; (right) median intensity—data plotted are mean ± SD; * p≤ 0.05 (One-way-ANOVA). **c**, (Left), MAP2 immunofluorescence staining of cortical neurons at day 40 generated with antioxidant (+AO) or without antioxidants (-AO+vA^lo^) or supplemented with ferrostatin-1 (-AO+vA^lo^+Fer-1). (Right), high-content image analyses (n=3); data plotted are mean ± SD. * p≤ 0.05 (One-way-ANOVA). **d**, MAP2 mRNA expression of cortical neurons generated with antioxidant (+AO) or without antioxidants (-AO+vA^lo^) or supplemented with ferrostatin-1 (-AO+vA^lo^+Fer-1) using quantitative RT-PCR (n=3). Data plotted are mean ± SD. ** p≤ 0.01, * p≤ 0.05 (One-way-ANOVA). **e**, Area of organoids at day 60 generated with antioxidant (+AO) or without antioxidants (-AO+vA^lo^) or supplemented with ferrostatin-1 (-AO+vA^lo^+Fer-1), and measured by ImageJ (n=6). Data plotted are mean ± SD. ** p≤ 0.01, * p≤ 0.05 (One-way-ANOVA). **f**, (Left), organoid sections stained for the ferroptosis marker TfR1 at day 60 of forebrain generation. (Right), measurement of the TfR1 fluorescence intensity using ImageJ. Data plotted are mean ± SD. * p≤ 0.05 (One-way-ANOVA); n=3 refers to biologically independent experiments.

We further investigated the role of ferroptosis in neuronal development in a more physiological context by using forebrain organoids. These organoids were generated using media containing +AO, -AO+vA^lo^, or -AO+vA^lo^+Fer-1. Interestingly, organoids without antioxidants (-AO+vA^lo^) exhibited a significantly reduced size at day-40—as demonstrated by the analysis of the area of the organoids— compared to +AO organoids, or organoids rescued with ferrostatin-1 (-AO+vA^lo^+Fer-1) (**Fig. 3e**). We recently demonstrated that the transferrin receptor (TfR1) can be utilized to visualize ferroptotic events^22,23^. Therefore, we sectioned organoids generated in +AO, -AO+vA^lo^ and -AO+vA^lo^+Fer-1 conditions at day 60 and stained these with antibodies against TfR1. Positive TfR1 staining could only be observed and quantified in -AO+vA^lo^ treated organoid sections, but was undetectable in +AO and -AO+vA^lo^+Fer-1 conditions, respectively (**Fig. 3f**), indicating that ferroptosis is present in conditions lacking antioxidants and can be eliminated when these organoids are treated with the ferroptosis-specific inhibitor ferrostatin-1. Together, the results show that omission of antioxidants during cortical neuronal differentiation increases ferroptotic cell death and that specific inhibition of ferroptosis is sufficient for cortical neuron differentiation.

### Ferroptosis inhibition is critical for proper laminar organization of cortical organoids

We next investigated the influence of ferroptosis on laminar organization of cortical organoids, a model of neurodevelopment with physiological relevance. At day 60, organoids generated without antioxidants (-AO+vA^lo^) resulted in a diffused and unorganized distribution of CTIP2+ (**Fig. 4a**), SATB2+ (**Fig. 4b**) and TBR1+ (Fig **4c**) neurons, all markers for proper patterning. In contrast, CTIP2+, SATB2+ and TBR1+ neurons in the organoids were structured in the presence of antioxidants (+AO) or ferrostatin-1 (-AO+vA^lo^+Fer-1), thereby generating a sharp CTIP2, TBR1 as well as SATB2 layer (**Fig. 4a, 4b, 4c**). Moreover, ferroptosis events in -AO+vA^lo^ organoids also altered the well-defined ventricular-zone-like structure in forebrain organoids, which contains SOX2+ neural progenitor cells (**Extended Data Fig. 5**).

**Fig. 4:**
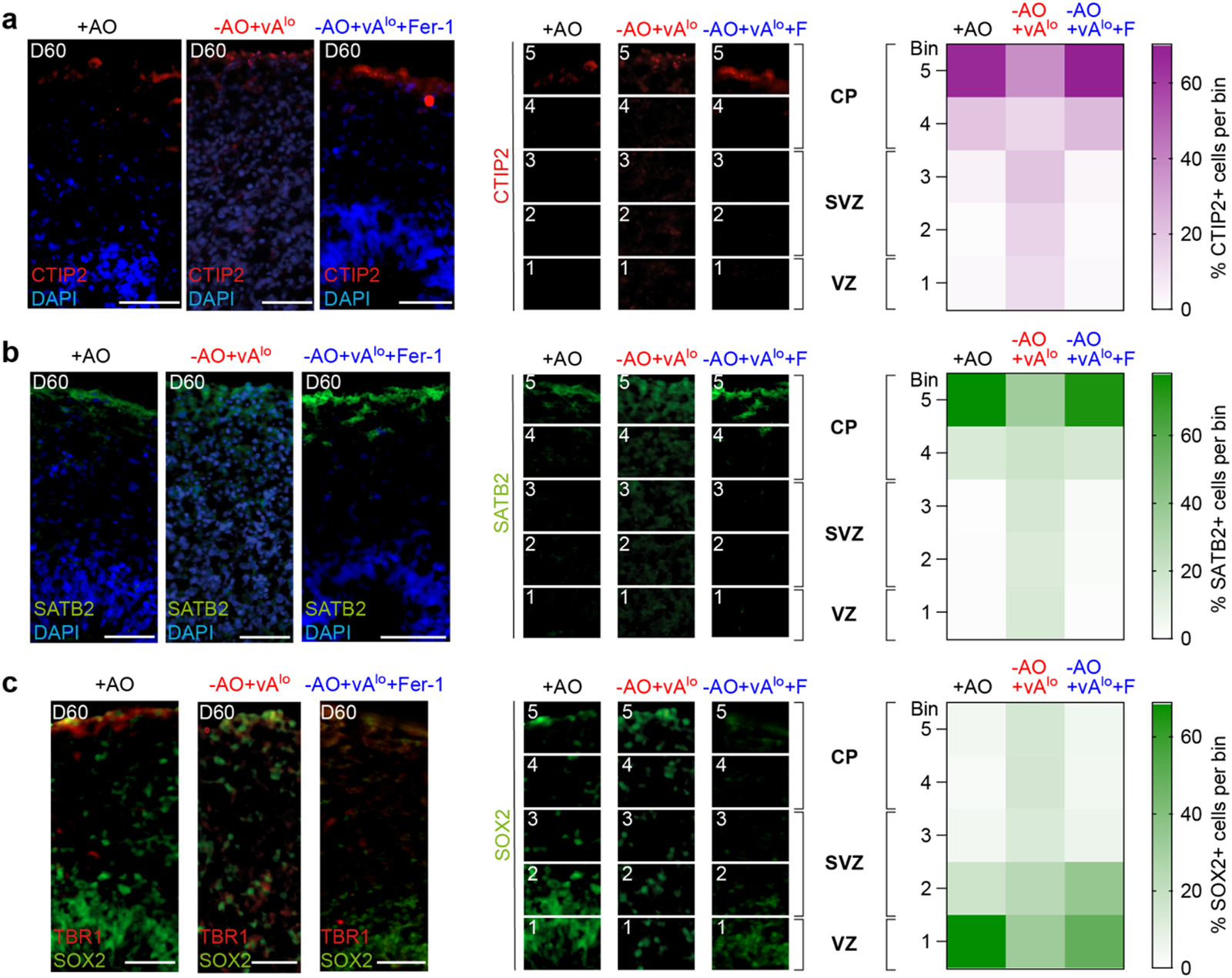
Inhibition of ferroptosis is critical for correct laminar organization of cortical organoids. **a-c**, (Left), immunofluorescence of day-60 forebrain organoid sections generated with antioxidant (+AO) or without antioxidants (-AO+vA^lo^), or the latter supplemented with ferrostatin-1 (-AO+vA^lo^+Fer-1), and stained for CTIP2/DAPI (**a**), SATB2/DAPI (**b**), and TBR1/SOX2 (**c**). Scale bars, 50 µm. (Middle), quantification of CTIP2+, SATB2+ and SOX2+ cells using ImageJ analysis. n = 3 biological replicates; * p < 0.05, ***p < 0.001, and ****p < 0.0001 (two-way ANOVA). CP, cortical plate; SVZ, subventricular zone; VZ, ventricular zone. (Right), distribution of CTIP2+, SATB2+ and SOX2+ cells illustrated using heatmaps; F = Fer-1

The distribution of CTIP2+, TBR1+, SATB2+ and SOX2+ cells was further quantified by dividing the developing cortex into five equal bins, as described previously^18,24^, with bin 1 corresponding to ventricular zone (VZ), bins 2 and 3 corresponding to subventricular zone (SVZ), and bins 4 and 5 corresponding to the cortical plate (CP) of the cortex (**Fig. 4a, 4b, 4c, Extended Data Fig. 5**). Cells without antioxidants (-AO+vA^lo^) undergoing ferroptosis showed significantly more CTIP2+ and SATB2+ cells localized in bins 1 to 4 and fewer cells localized in bin 5 compared to control cells treated with antioxidants (+AO) or Fer-1 (-AO+vA^lo^+Fer-1) (**Fig. 4a, 4b**). The inverse distribution was observed for TBR1+ neurons; *i.e.,* more cells in bins 3 to 5 in conditions without antioxidants (-AO+vA^lo^) compared to control cells (+AO) or Fer-1 treated cells (-AO+vA^lo^+Fer-1) (**Fig. 4c**). The same observation was made for SOX2+ cells, which were shifted more towards bins 3 to 5 in antioxidant-negative conditions (**Extended Data Fig. 5**). Hence, our data demonstrate impaired laminar organization of cortical organoids when organoids were grown without antioxidants, and this phenotype could be reversed when ferroptosis was specifically inhibited with ferrostatin-1, proving the importance to suppress ferroptosis during neuronal differentiation.

## Discussion

In this study, we found that overcoming ferroptosis is a critical requirement for the differentiation process of pluripotent stem cells into neurons. We demonstrate that ferroptosis can be inhibited by the use of either antioxidants (GSH and vitamin E) or ferrostatin-1, and we have identified vitamin A as a novel ferroptosis suppressor (model in **Extended Data Fig. 6**). Interestingly, protocols for stem cell differentiation mostly contain antioxidants^16,25–30^, but the exact mechanism of why these antioxidants are essential has been unclear. The differentiation process uses transferrin and PUFAs (Supplementary Table 1), both of which are involved in promoting lipid peroxidation. *In vivo*, ROS levels are important for proliferation and differentiation events during early mouse embryonic brain development, but continuous increase of ROS leads to oxidative stress and lipid peroxidation^31^. Therefore, a tight control is needed. Our work resolves that during early phases of cortical neuron differentiation antioxidants prevent cells from lipid peroxidation and ferroptosis to enable neuronal development. Notably, ferroptotic cell death could affect neural progenitor (radial glia) cells and/or the maturing neurons. From our organoid data we assume that progenitor cells undergo ferroptosis as less TBR1+ cells reach the upper layer (cortical plate) leading to enrichment of cells in the VZ and SVZ. Future research will decipher the exact cell types affected by ferroptosis during neurogenesis. Physiologically, a recent study showed that vitamin E is necessary for zebrafish nervous system development^32^. Also, vitamin E deficiency leads to accumulation of lipid peroxides in the zebrafish nervous system and impairs cognitive functions^33^, which supports our findings. Notably, our discovery of inhibiting ferroptosis during early neuronal development may be generalizable to developmental processes: ROS levels are increased during mammalian reproduction from fertilized egg to the implantation, where ROS is needed for signaling purposes and embryo development; however, elevated levels of ROS can be lethal^34^. Therefore, the female reproductive tract is enriched with vitamin E, vitamin A and GSH^34^ that are able to directly or indirectly act as antioxidants to control ROS levels.

Vitamin A and its active metabolite retinoic acid (all-trans retinoic acid, ATRA) are known to have major impact on embryonic development including development of the brain, limbs, eyes, the heart, skeleton, and more^35^. This influence is based on the ability to activate the nuclear receptors RAR/RXR for transcriptional induction of important developmental genes^4^. In mice, retinoic acid is expressed very early during neuronal development and activates a series of important genes that control brain patterning and progenitor differentiation^35^. At the same time, there is evidence that vitamin A has antioxidant capacity^36^. While there has been long-lasting debate about a direct or indirect antioxidant effect of vitamin A, there is now a consensus understanding that vitamin A indirectly acts as an antioxidant by transcriptionally regulating antioxidative mediators^36^. This is in line with our data, as we are unable to detect any direct radical-trapping activity of ATRA in cell-free C11-BODIPY and DPPH assays. In this study, we unravel that vitamin A activates RAR/RXR to upregulate GPX4, FSP1, GCH1, ACSL3, among other genes that are established as key gatekeepers to eliminate lipid hydroperoxides and to inhibit ferroptosis. This mechanism is distinct from vitamins E and K to counteract ferroptosis^8,11,12^, which act through radical-scavenging activity. Vitamin A at concentrations capable of upregulating GPX4, FSP1, GCH1, and ACSL3 is sufficient to overcome ferroptotic cell death during early neurogenesis and facilitate neuronal development. Together, our study answers a substantial question of why antioxidants are needed in early neurogenesis and establishes vitamin A as a potent ferroptosis inhibitor with a distinct antioxidant mechanism.

## Methods

### Chemicals

For induction of ferroptotic cell death we used (1S,3R)-RSL3 (Sigma-Aldrich) and Imidazole Ketone Erastin (IKE, Cayman Chemical). As ferroptosis inhibitors we used ferrostatin-1 (Fer-1, Sigma-Aldrich) and all-trans-Retinoic acid (ATRA, Sigma-Aldrich).

For inhibition of different receptors and proteins we used HX 531 (Biomol) as a pan-Retinoic Acid Receptor (RAR) antagonist, AGN 193109 (Sigma-Aldrich) as a Retinoic X Receptor (RXR) antagonist, iFSP1 (MedChemExpress) as an inhibitor of Ferroptosis Suppressor Protein 1 (FSP1) and GW6471 (Selleck Chemicals) as a PPARα antagonist.

### Cell culture

#### HT-1080

The fibrosarcoma cell line HT-1080 (purchased from ATCC) was cultured in Dulbecco’s Modified Eagle’s Medium supplemented with 10% fetal bovine serum, 1% non-essential amino acids (MEM NEAA) and 1% Penicillin-Streptomycin (all purchased from Thermo Fisher Scientific). Cell line was grown at 37°C and 5% CO_2_.

#### Human stem cells

H9 (WA09) cells were maintained with Essential 8 Flex media (#A28558501, Thermo Fisher Scientific) in feeder-free conditions on vitronectin (VTN-N) (#A14700, Thermo Fisher Scientific). H9 (WA09) cells were passaged as clumps with 0.5M EDTA (0.5M EDTA, 5M NaCl, 1X PBS). The pluripotency level of the cells in culture was routinely authenticated for the markers Nanog and Oct-4 and checked for mycoplasma contamination.

### Cell viability assay

To test cell viability or create dose-response curves, 750 HT-1080 cells per well were seeded into 384-well plates (CulturPlate, PerkinElmer) and incubated for 24 h. Cells were treated with indicated RSL3 concentrations and 20 µM ATRA for 18 h; for dose-response curves, compounds were diluted in a 12-point series. Experiments with small molecule inhibitors were performed with the following concentrations: 200 nM RSL3, 1,5 µM IKE, 20 µM ATRA and indicated concentrations of RARi, RXRi, iFSP1 or PPAR*α*i, respectively. After treatment for 18 h, 20 µl CellTiter-Glo 2.0 Reagent (Promega) was added directly into each well and luminescence was detected using an EnVision 2104 Multilabel plate reader (PerkinElmer).

### Spheroid formation, treatment and imaging

Spheroids were cultured according to the protocol previously described^15^. In brief, HT-1080 were seeded into 96-well round bottom microplates (CellCarrier Spheroid ULA, PerkinElmer) with a density of 2,000 cells per well. After 48 hours of proliferation, spheroids were treated with 200 nM RSL3, 20 µM ATRA, 2 µM Fer-1, 10 µM RARi and 10 µM RXRi, and incubated another 48 h. Finally, spheroids were stained with a 1:10,000 dilution of Hoechst 33342 (Sigma) and after 1 h of incubation at 37 °C, imaging was performed using an Operetta high-content system. For image analysis, the Columbus software (PerkinElmer) was used.

### Antibody staining and flow cytometry

Immunostaining of HT-1080 with anti-4-Hydroxynonenal-antibody (4-HNE, ab46545, Abcam) was performed as previously described^15^. In brief, ferroptotic cell death was induced via 300 nM RSL3 for 2 h and cells were co-treated with 20 µM ATRA or 2 µM ferrostatin-1. 10% normal goat serum (Thermo Fisher Scientific) was used for blocking before cells were incubated in anti-4-HNE antibody. As a secondary antibody, anti-rabbit Alexa 488 antibody (A32731, Thermo Fisher Scientific) was used. For flow cytometry, 10,000 events per condition were measured in an Attune acoustic flow cytometer (Applied Biosystems). FlowJo v10.8.1 Software (BD Life Sciences) was used to create histograms and analyze intensities.

### BODIPY C11 staining

#### Microscopy

HT-1080 cells were seeded into 6-well plates with a density of 400,000 cells per well and incubated for 24 h, then 1 µM C11-BODIPY 581/591 (Thermo Fisher) was added for 30 min. Without changing the medium, cells were then treated with 250 nM RSL3 and 20 µM ATRA. After 2 h of treatment, cell culture medium was replaced with PBS and fluorescence was detected in the GFP channel of an EVOS FL fluorescence microscope (Thermo Fisher Scientific) using 20-fold magnification.

#### Flow cytometry

C11-BODIPY staining and flow cytometry was performed as previously described^15^. Ferroptosis was induced for 2 h with 250 nM RSL3 and cells were co-treated with 20 µM ATRA or 2 µM ferrostatin-1. HT-1080 cells were stained with 2 µM C11-BODIPY for 30 min. Day-20 neurons were stained with 2 µM C11-BODIPY for 1 h. Afterwards cells were measured in an Attune acoustic flow cytometer (Applied Biosystems). FlowJo v10.8.1 Software (BD Life Sciences) was used to create histograms and analyze intensities.

### TBARS Assay

HT-1080 cells were seeded into 100 mm dishes with a density of 2.5 million cells per dish and incubated for 48 h. Cells were then treated with 250 nM RSL3 for 3 h and co-treated with 20 µM ATRA or 2µM ferrostatin-1 before they were harvested using 0.05% Trypsin-EDTA (Thermo Fisher Scientific) and cell scraper (Sarstedt). Cell counts of every treatment condition were determined and adjusted to the condition with the lowest count. TBARS (TCA Method) Assay Kit (Cayman Chemical, 700870) was performed according to manufacturer’s instructions to detect levels of Malondialdehyde (MDA). Because the fluorometric method was chosen for detection, sample fluorescence was measured at 530 nm / 550 nm in an EnVision 2104 Multilabel plate reader (PerkinElmer).

### RNA isolation and quantitative RT-PCR

#### HT-1080

After treatment of HT-1080 cells with 20 µM ATRA for 7h, total RNA isolation was performed by using the Monarch Total RNA Miniprep Kit (New England BioLabs) and RNA concentration was detected in a NanoDrop 2000 spectrophotometer (Thermo Fisher Scientific). For synthesis of complementary DNA, we used the Maxima H Minus First Strand cDNA Synthesis Kit including DNA digestion (Thermo Fisher Scientific) and oligo (dT)_18_ primer and random hexamer primer. Quantitative RT-PCR was performed with PowerUp SYBR Green Master Mix (Thermo Fisher Scientific) in a LightCycler480 (Roche) using the respective primers listed in Table 1. Quantification of gene expression was done using the 2∧(-*ΔΔ*Ct) method^37^ and obtained expression levels were normalized to GAPDH expression.

#### Cortical neurons

Total RNA from cortical neurons was isolated using TRIzoL (#15596-026, Invitrogen) following manufacturer’s protocol. RNA extraction was performed using chloroform. RNA was precipitated in isopropanol and resuspended in nuclease free water. Total RNA (500 ng) was reverse transcribed using the Superscript First Strand Synthesis kit following manufacturer’s protocol and random hexamers after DNAse treatment. qRT-PCR was performed using specific primers and the LightCycler 480 SYBR Green I (#04887352001, Roche) on a LightCycler 480 (Roche). Relative gene expression was calculated using the 2∧(-ΔCt) method^38^. All genes were normalized to RNA polymerase II values.

Primer details are provided in Table 1.

**Table 1:**
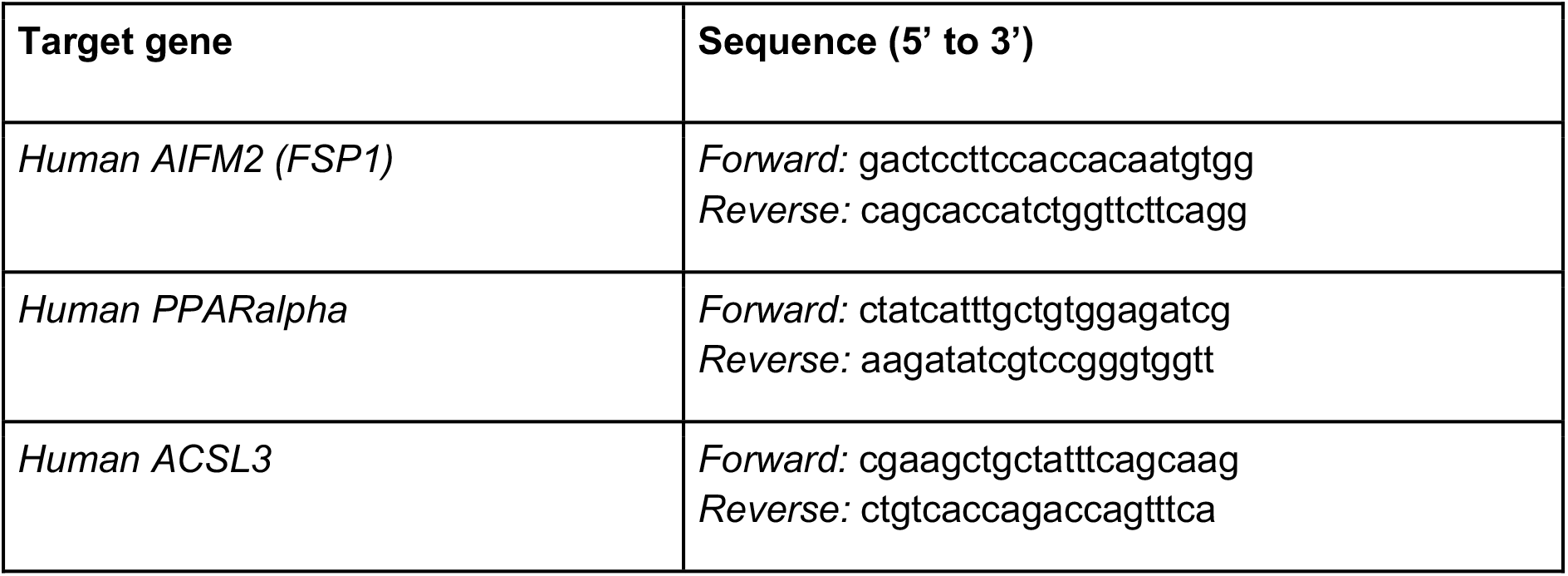
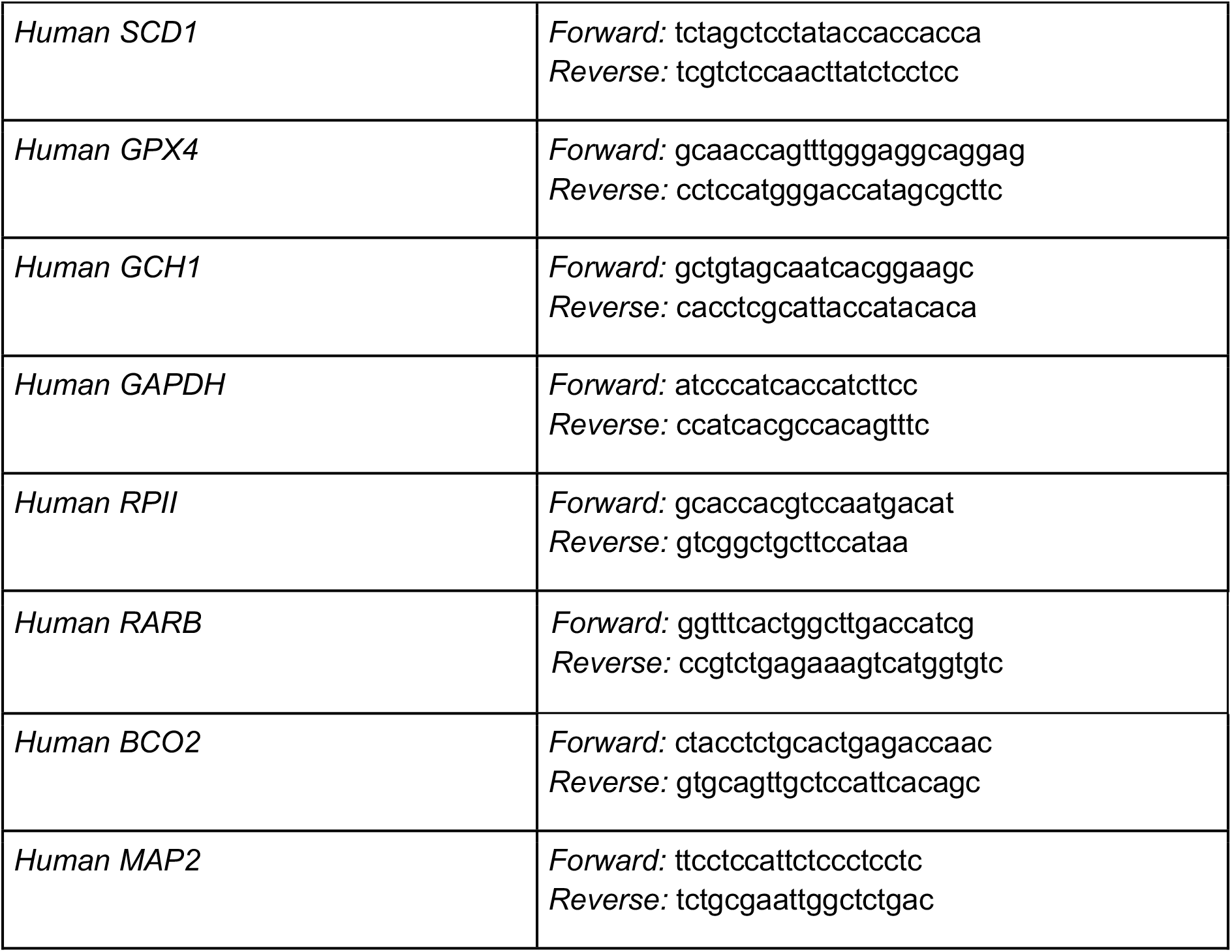
qRT-PCR primers.

### Cell-free C11-BODIPY and DPPH Assays

For the cell-free C11-BODIPY assay, ATRA was diluted in 150 µl PBS to a final concentration of 25 µM. C11-BODIPY 581/591 (Thermo Fisher Scientific) was diluted to 1.875 µM and 2,2’-Azobis(2-methylpropionamidine) dihydrochloride (AAPH, Sigma) was diluted to 7.5 mM, each in separate tubes of 150 µl PBS. All three tubes were mixed, vortexed and incubated for 30 minutes at room temperature and protected from light. Triplicates of 100 µl were transferred into a black 96-well plate (Greiner Bio-One) and fluorescence (495nm/520nm) was measured with an EnVision 2104 Multilabel plate reader (PerkinElmer). As control samples, the same procedure was performed with ferrostatin-1 (25 µM) as a positive control, DMSO as a negative control and one sample without the radical-inducing AAPH.

The 2,2-diphenyl-1-picrylhydrazyl (DPPH) Assay was performed according to a published protocol^8^. In brief, ATRA was diluted in DMSO to a final concentration of 10 mM. 5 µl of this dilution were added to 1 ml of 0.05 mM DPPH (Sigma, D9132) in methanol and rotated at room temperature for 10 min. 200 µl were transferred into a clear bottom 96-well plate (Corning costar) in quadruplicates and absorbance was measured at 517 nm with an EnVision 2104 Multilabel plate reader (PerkinElmer). As control samples, ferrostatin-1 and DMSO were measured with the same procedure. Background absorbance of methanol-only was subtracted from measured values.

### Differentiation of stem cells into cortical neurons

#### Coating plates for differentiation

Prior to the start of differentiation experiments, 24-well cell culture plates were coated with Geltrex (#A14133-02, Thermo Fisher Scientific) as previously described^18^. Briefly, Geltrex was thawed overnight on ice at 4°C. The next day Geltrex was diluted in ice cold DMEM/F12 (#11330-032/ Thermo Fisher Scientific) (ratio 1:30) and added to the plates (0.5 ml/well for 24-well plate or 2 ml/well for 6-well plate). Plates were sealed with parafilm and stored overnight at 4°C.

#### Coating plates for replating with PO/Lam/FN

Before replating, poly-L-ornithine hydrobromide (PO) (#P3655/Sigma) was diluted to 15 mg/ml in PBS. 0.5 ml/well for 24-well plate was added and incubated overnight at 37°C / 5% CO2. Next day, the PO solution was removed and the plates were washed 3-times with 1x PBS. Afterwards Laminin I (3400-010-1, R&D systems) and Fibronectin (FC010, EMD Millipore Corp.) were diluted at 2mg/ml in PBS and added at 0.5 ml/well to a 24-well plate. The plates were incubated overnight at 37°C.

#### Differentiation into cortical neurons

The cortical neuron differentiations were carried out as previously described^16-18^. Briefly, human pluripotent stem cells were dissociated using Accutase and plated at high density on Geltrex (#A14133-02, Thermo Fisher Scientific) coated wells. Next day, the plates were checked for complete confluency and directed towards cortical neuron patterning. Cells were cultured in Essential 6 medium (E6, #A1516401, Thermo Fisher Scientific) in the presence of LDN193189 (#72142, Stem Cell Technologies), SB431542 (#1614, Tocris) and XAV939 (#3748, Tocris) (until day 5). From day 5 to 10, cells were cultured in the presence of TGF*β* and BMP inhibitors (LDN193189, #72142, Stem Cell Technologies; SB431542, #1614, Tocris) to trigger cortical precursors for cortical neuron differentiation. From day 10 to 20, the cells were cultured in Neurobasal media (#21103-049, ThermoFisher Scientific) and DMEM F-12 media (#11330-032/ Thermo Fisher Scientific) supplemented with 1% Pen/Strep (#15140-122, Thermo Fisher Scientific), L-Glutamine (#25030-024, Thermo Fisher Scientific), B27(-vitamin A) (#12587010, Thermo Fisher Scientific), N2 supplement B (# 07156, StemCell Technologies), D-(+) Glucose, 2-mercaptoethanol, Sodium bicarbonate and Progesterone. From day 20 on cortical neurons were cultured on polyornithine (PO)/ laminin (L)/fibronectin (FN) coated wells and kept in neuronal differentiation media (Neurobasal media (#21103-049, Thermo Fisher Scientific), 1% Pen/Strep (#15140-122, Thermo Fisher Scientific), L-Glutamine, B27(-vitamin A) supplemented with DAPT (#2634, Tocris), BDNF (#450-10, PeproTech), GDNF (#248-BD-025, R&D biosystems), cAMP (#D0627, Sigma) and AA (#4034-100, Sigma)) as described previously^16,18,28,39.^

### Immunofluorescence of cortical neurons

At day 20, neurons were washed with 1X PBS and fixed with 4% PFA for 15 min at room temperature. Cells were permeabilized with 0.03% Triton X-100/PBS for 10 min and washed twice in 0.15% Triton X-100/PBS at room temperature. Blocking was done in 0.15% Triton X-100/PBS supplied with 5% BSA (40mg/mL) for 60 minutes at room temperature. Cells were incubated with the respective primary antibody (Mouse monoclonal Anti-MAP2 Sigma-Aldrich M-1406, RRID:AB_477171) in 0.15% Triton X-100/PBS supplied with 5% BSA (40mg/mL) overnight at 4°C. Next day, cells were washed three times with 0.15% Triton X-100/PBS and incubated with the respective secondary antibody (AlexaFluor Goat Anti-Mouse 488 Thermo Fischer Scientific R37120, RRID:AB_2556548) in 0.15% Triton X-100/PBS supplied with 5% BSA for 2 h. Nuclear stain DAPI was added at a concentration of 1:10,000 90 minutes post incubation with secondary antibody. Finally, cells were washed three times with 0.15% Triton X-100/PBS and imaged using a Zeiss (AX10) or Zeiss LSM980 microscope.

### Generation of forebrain organoids

Forebrain organoids were generated from human pluripotent stem cells as described in^18,40^. Briefly, H9 cells were dissociated in Accutase. 9,000 cells/well were seeded in a V-bottom ultra-low adhesion 96 well plate (#MS-9096VZ, Sbio prime surface 96V plate) in Essential 8 flex medium + Y-drug (#1254 Y-27632 dihydrochloride, Tocris), centrifuged at 3,000 rpm for 10 minutes. From D0-D5, cells were cultured in Essential 6 medium (E6, #A1516401, Thermo Fisher Scientific) in the presence of LDN193189 (#04-0074, Stemgent), SB431542 (#1614, Tocris) and XAV-939 (#3748, Tocris). XAV-939 was removed from day 4 till day 18. From D18 onwards, organoids were maintained in organoid differentiation medium on an orbital shaker (50% DMEM F-12 (#11320-033, Thermo Fisher Scientific), 50% Neurobasal media (#21103-049, Thermo Fisher Scientific), 0.5x N2 supplement (#17502-048, Thermo Fisher Scientific), 0.025% insulin (#I-034, Sigma), 5 mM L-Glutamine (#25030024, Thermo Fisher Scientific), 0.7 mM MEM-NEAA (#11140050, Thermo Fisher Scientific), 50 U/mL Penicillin-Streptomycin (#15140-122, Thermo Fisher Scientific), 55 mM 2-mercaptoethanol (#21985-023, Thermo Fisher Scientific), 1xB27 supplement -vitamin A (#12587010, Thermo Fisher Scientific).

### Sectioning and immunostaining of forebrain organoids

For visualizing organoids, cells were fixed in 4% PFA overnight at 4°C and washed three times in 1X PBS. After fixation, organoids were cryoprotected in 30% sucrose/PBS on a rotor shaker overnight at 4°C and sectioned at 20 µm on a cryostat (Leica 1850 UV). Sections were permeabilized in 0,3% Triton X-100/PBS, blocked in 0.15% Triton X-100/PBS supplied with 5% BSA (40 mg/mL) and incubated as floating sections in primary antibody (Mouse monoclonal Anti-SOX2 SantaCruz Biotech sc-365823, RRID:AB_10842165; Rat monoclonal Anti-CTIP2 Abcam ab18465, RRID:AB_2064130; Rabbit polyclonal Anti-TBR1 Abcam ab31940, RRID:AB_2200219; Rabbit polyclonal Anti-SATB2 Sigma-Aldrich HPA001042, RRID:AB_10601711; Mouse Anti-TfR1 (CD71—3B8 2A1) antibody Santa Cruz sc-32272) overnight. The next day, sections were washed three times with PBS and incubated with secondary antibody (AlexaFluor Goat Anti-Mouse 488 Thermo Fischer Scientific R37120, RRID:AB_2556548; AlexaFluor Goat Anti-Rabbit 488 Thermo Fischer Scientific A32731, RRID:AB_2633280; AlexaFluor Goat Anti-Rat 488 Abcam ab150157, RRID:AB_2722511; AlexaFluor Goat Anti-Mouse 594 Abcam ab150116, RRID:AB_2650601; AlexaFluor Goat Anti-Rabbit 594 Thermo Fischer Scientific A11012, RRID:AB_141359; AlexaFluor Goat Anti-Rat 568 Thermo Fischer Scientific A-11077, RRID:AB_141874; AlexaFluor Goat Anti-Rabbit 647 Thermo Fischer Scientific A-27040, RRID:AB_2536101) for 2 hours at room temperature. Nuclear stain, DAPI was added at a concentration of 1:10,000 90 minutes post incubation with the secondary antibody. Afterwards, sections were mounted onto microscopic slides using immu-mount, covered with coverslips and imaged using a Zeiss (AX10) or Zeiss LSM980 microscope.

### Treatment of cortical neurons and forebrain organoids

Between day 10 and 20 during cortical differentiation B27 supplement was replaced with B27 supplement without antioxidants (B27 supplement-AO, #10889038, Thermo Fisher Scientific) and B27 supplement without antioxidants + 10 µM all-trans Retinoic Acid (ATRA) or 1 µM ferrostatin-1, respectively. During the forebrain organoid generation organoids were kept in organoid differentiation media without antioxidants (B27 supplement-AO, #10889038, Thermo Fisher Scientific) and 1 µM ferrostatin-1, respectively, from day 18 onwards.

### Image analysis

High-content image analysis of neurons was performed as previously described^18^. For image analysis of spheroids and BODIPY staining, Columbus software (PerkinElmer) was used. Spheroids were detected as “Image region” and morphology properties were calculated (*e.g.,* roundness). For BODIPY staining, the number of green fluorescent objects per well was counted.

### Statistical analysis

All statistical testing for significance was performed in GraphPad Prism Software version 9.4. Differential gene expression analysis was conducted in R software version 4.2.0 using the DESeq2 package^41^. Statistical details to every experiment are described in the figure legends.

### Code availability

Source code is available at https://github.com/MendenLab/DEx_StemCells_new.

## Acknowledgement

We thank Lena Klepper and Stefanie Brandner for excellent technical assistance.

## Author contributions

K.H. and M.V. conceptualized, designed and generally supervised the study; J.T., I.R., and K.H. performed and analyzed the HT-1080 experiments; V.P.N., G.C., J.Tch., H-M.T., M.V. performed, analyzed and interpreted the stem cell experiments; A.G., R.P., M.P.M. analyzed and interpreted the RNAseq data; B.R.S., L.S., M.P.M., M.V., K.H. supervised individual experiments; J.T., V.P.N., M.V., K.H. wrote the original draft; J.T., V.P.N., M.V., K.H. edited the manuscript; all authors read, commented and approved the manuscript for submission.

---

**Extended Data Fig 1.**
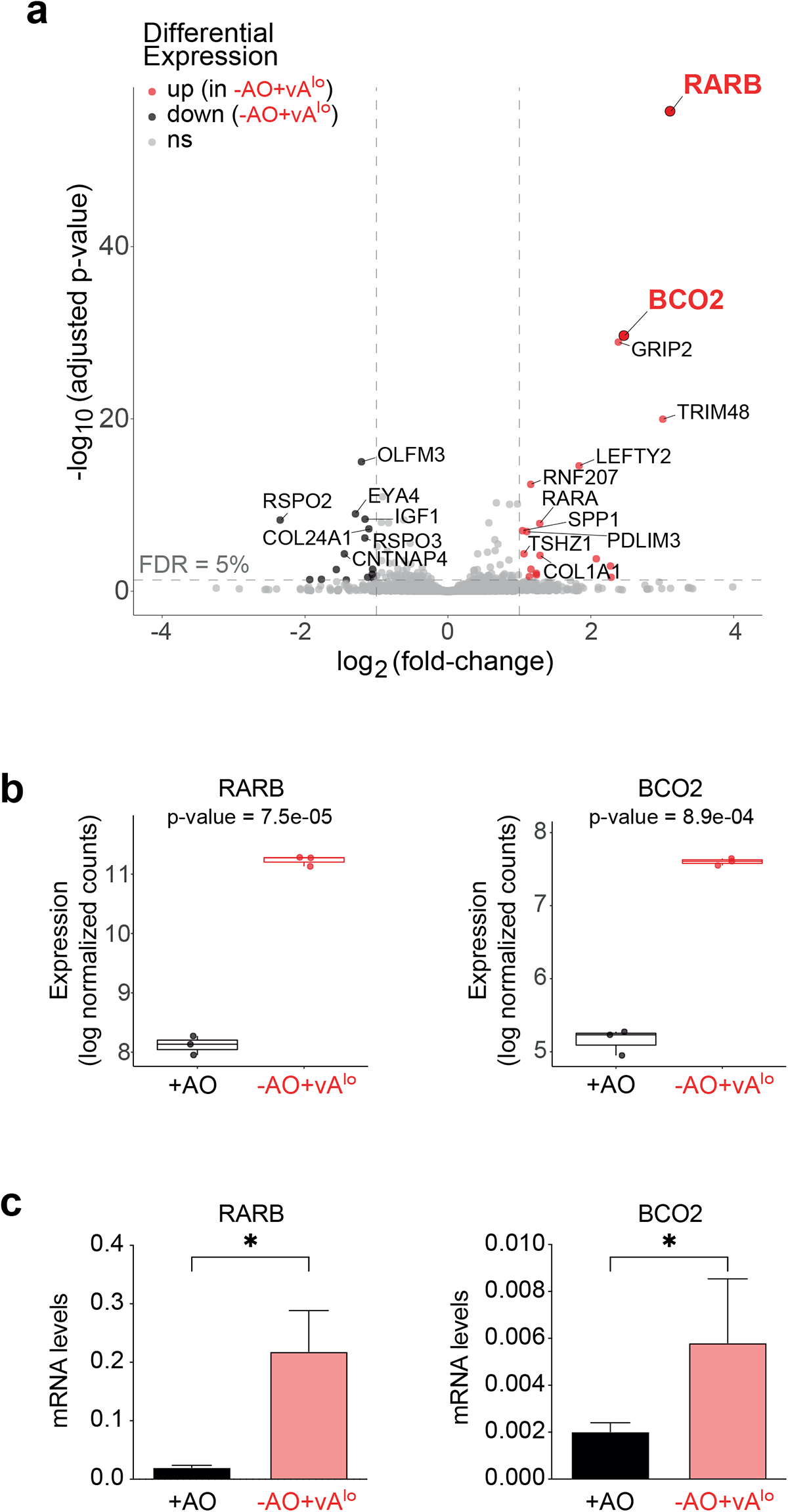
Vitamin A supplementation during cortical neuronal differentiation results in overexpression of Retinoic Acid Receptor B and beta-carotene oxygenase 2. **a**, Volcano plot of total RNA sequencing (RNA-seq) from cortical neurons (day-20) generated with antioxidants (+AO) or without antioxidants (-AO+vA^lo^). **b**, Box plot depicting expression of Retinoic Acid Receptor B (RARB) and beta-carotene oxygenase 2 (BCO2) in total RNA-seq of immature cortical neurons. **c**, Validation using quantitative RT-PCR (n=3). Data plotted are mean ± SD. ** p≤ 0.01, * p≤ 0.05 (unpaired t-test).

**Extended Data Fig 2.**
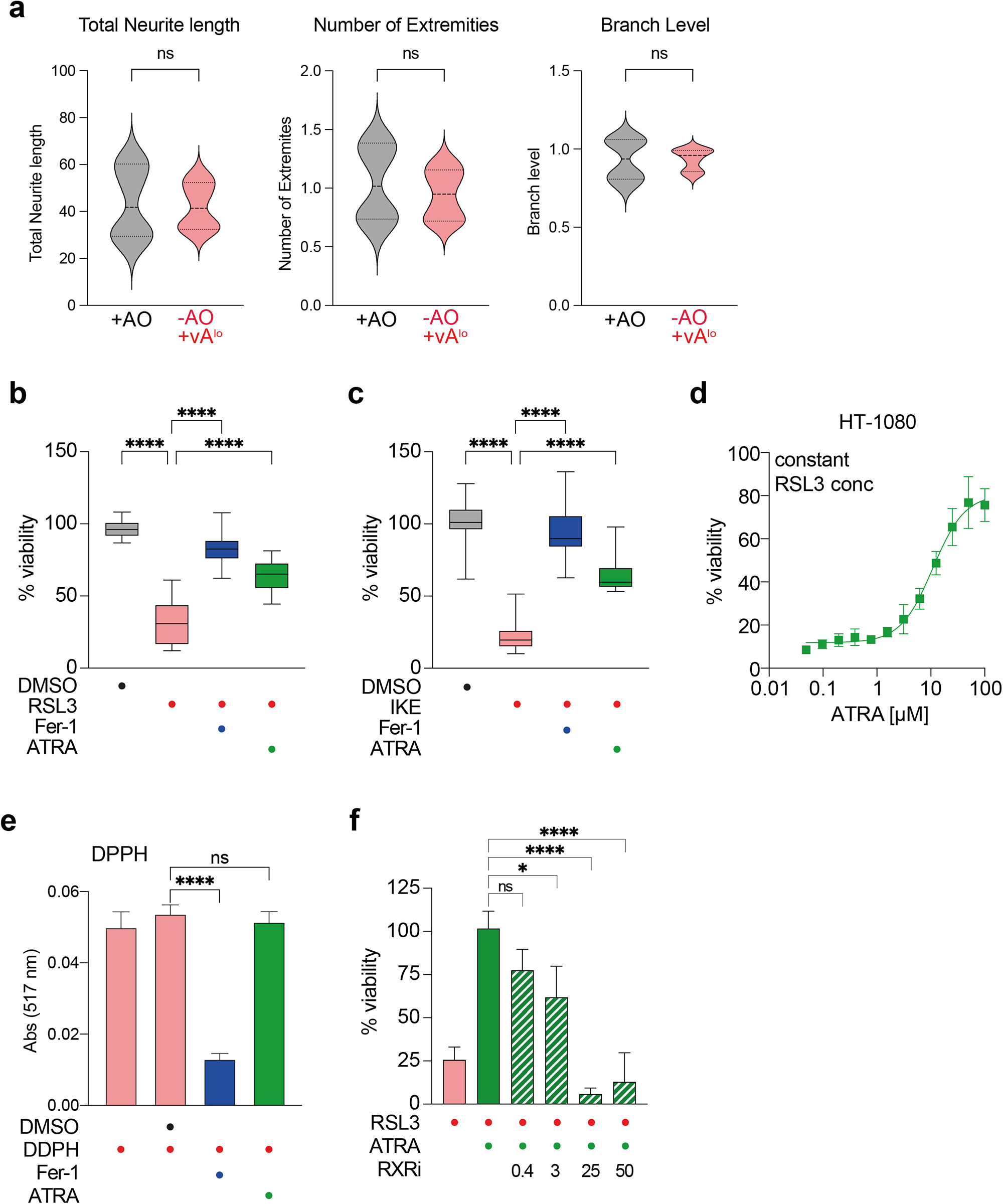
Effects of antioxidants and vitamin A on ferroptosis and lipid peroxidation. **a**, Quantification of morphological changes of cortical neurons upon differentiation with (+AO) or without antioxidants (-AO+vA^lo^)) using high-content Image analysis (n=3). **b**, CellTiter-Glo assay of HT-1080 cells co-treated with the ferroptosis inducer RSL3 and ferrostatin-1 (Fer-1) or vitamin A (ATRA) (n=3). Data plotted are mean ± SD. **** p≤ 0.0001 (One-way-ANOVA). **c**, CellTiter-Glo assay of HT-1080 cells co-treated with the ferroptosis inducer IKE and ferrostatin-1 (Fer-1) or vitamin A (ATRA) (n=3). Data plotted are mean ± SD. **** p≤ 0.0001 (One-way-ANOVA). **d**, CellTiter-Glo assay of HT-1080 cells co-treated with the ferroptosis inducer RSL3 and dose-response of vitamin A (ATRA). **e**, Absorbance of radical initiator 2,2-diphenyl-1-picrylhydrazyl (DPPH) co-treated with ferrostatin-1 (Fer-1) or vitamin A (ATRA) (n=3). Data plotted are mean ± SD. **** p≤ 0.0001 (One-way-ANOVA). **f**, CellTiter-Glo assay of HT-1080 cells after co-treatment with RSL3 and vitamin A (ATRA) as well as different concentrations of the small molecule inhibitor of Retinoid X receptor (RXRi) (n=3). Data plotted are mean ± SD. **** p≤ 0.0001, * p≤ 0.05 (One-way-ANOVA); n=3 refers to biologically independent experiments.

**Extended Data Fig 3.**
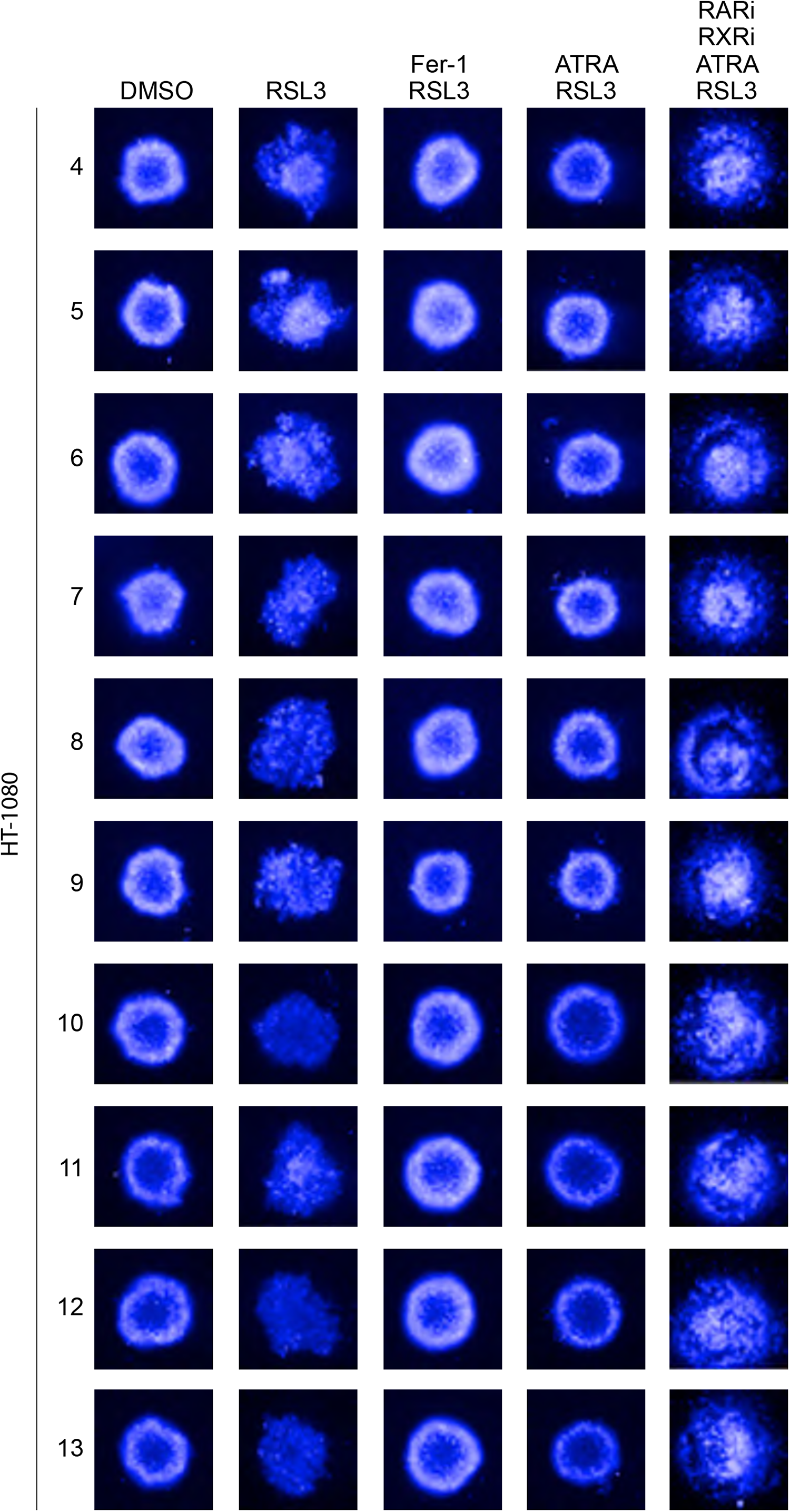
Inhibition of ferroptosis by vitamin A impacts spheroid growth. Remaining replicates of spheroids with the indicated treatments corresponding to Fig. 1h.

**Extended Data Fig 4.**
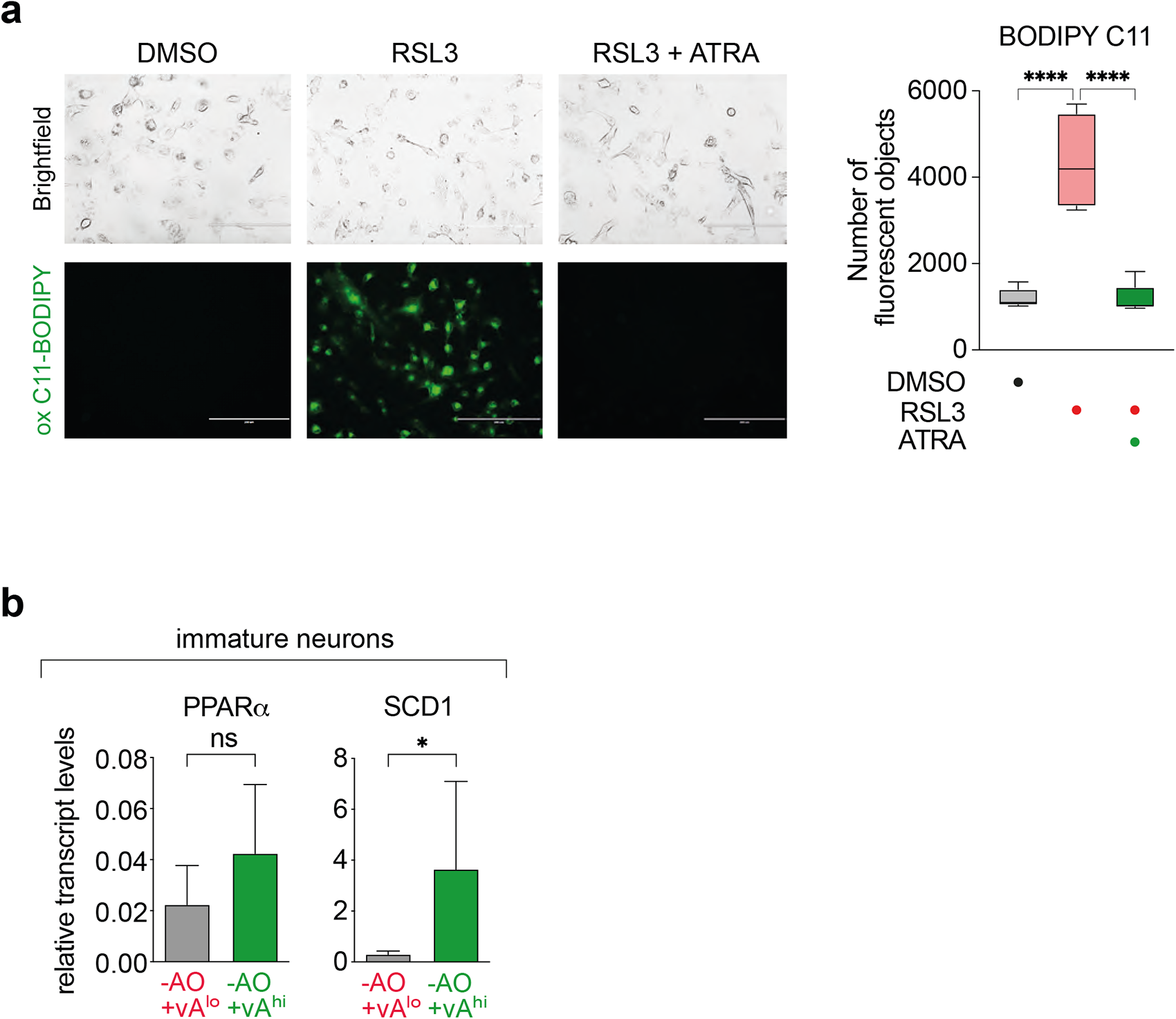
Vitamin A inhibits lipid peroxidation and affects ferroptosis target genes. **a**, (Left), C11-BODIPY microscopy images of HT-1080 cells co-treated with RSL3 and vitamin A (ATRA) (n=3). (Right), Quantification of fluorescent objects. **** p≤ 0.0001 **b**, Relative mRNA levels of PPAR*α* and SCD1 in immature neurons (day-20) measured by quantitative RT-PCR (n=3). Data plotted are mean ± SD. * p≤ 0.05, ns = not significant (unpaired t-test); n=3 refers to biologically independent experiments.

**Extended Data Fig 5.**
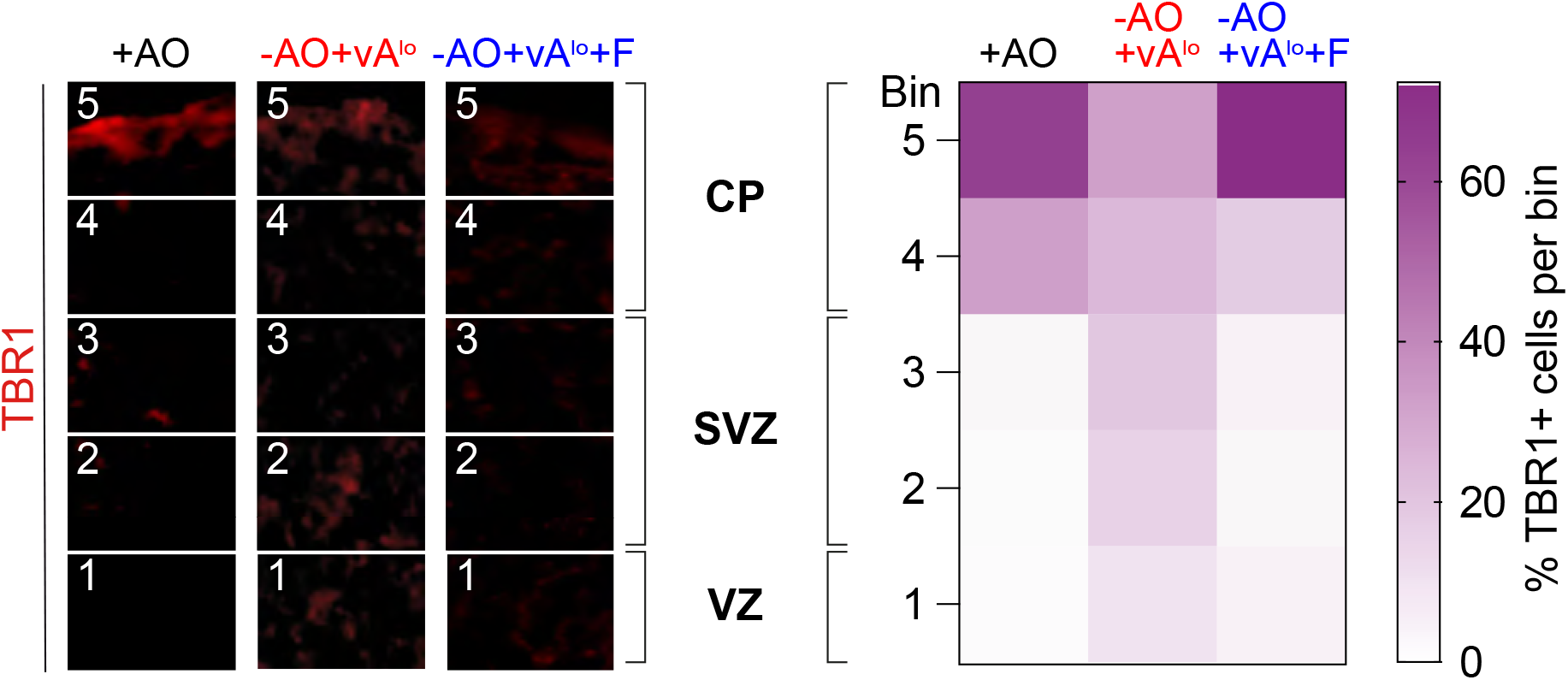
Inhibition of ferroptosis is critical for correct laminar organization of cortical organoids. **a-c**, Left, immunofluorescence of day-60 forebrain organoid sections generated with antioxidant (+AO) or without antioxidants (-AO+vA^lo^), or the latter supplemented with ferrostatin-1 (-AO+vA^lo^+Fer-1), and stained for TBR1. Scale bars, 50 µm. (Middle), quantification of TBR1+ cells using ImageJ analysis. n = 3 biological replicates; *p < 0.05, ***p < 0.001, and ****p < 0.0001 (two-way ANOVA). CP, cortical plate; SVZ, subventricular zone; VZ, ventricular zone. (Right), distribution of TBR1+ cells illustrated using heatmaps; F = Fer-1

**Extended Data Fig 6.**
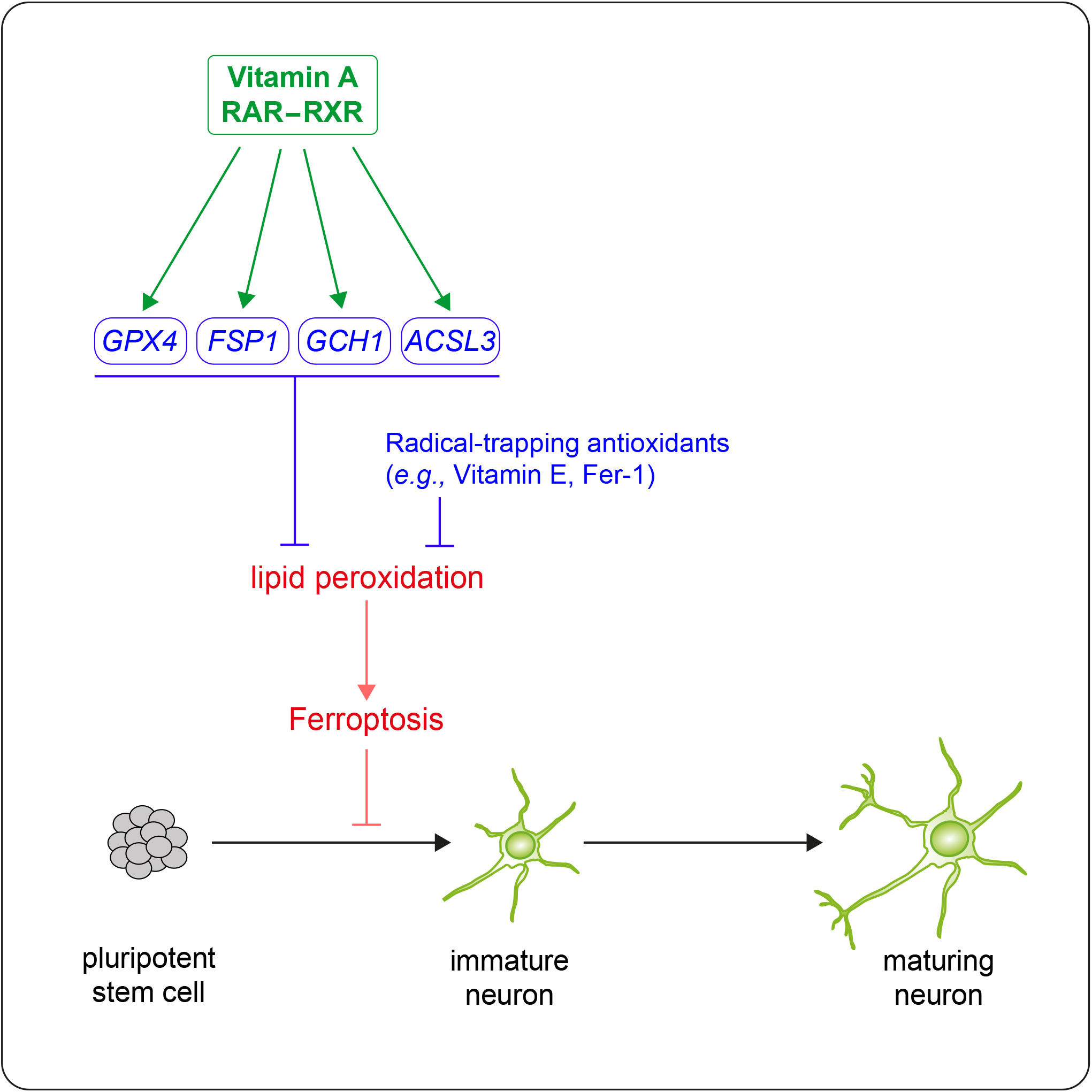
Model of vitamin A suppressing ferroptosis to promote neuronal development.

Ingredients according to “Media Formulations” on ThermoFisher Scientific Webpage

|  | **B27** | **B27(-AO+VA)** |
| --- | --- | --- |
| 1 | **Biotin** | **Biotin** |
| 2 | **DL Alpha Tocopherol Acetate** | N/A |
| 3 | **DL Alpha-Tocopherol** | N/A |
| 4 | N/A | **Vitamin A** |
| 5 | **BSA, fatty acid free Fraction V** | **BSA, fatty acid free Fraction V** |
| 6 | **Catalase** | N/A |
| 7 | **Human Recombinant Insulin** | **Human Recombinant Insulin** |
| 8 | **Human Transferrin** | **Human Transferrin** |
| 9 | **Superoxide Dismutase** | N/A |
| 10 | **Corticosterone** | **Corticosterone** |
| 11 | **D-Galactose** | **D-Galactose** |
| 12 | **Ethanolamine HCl** | **Ethanolamine HCl** |
| 13 | **Glutathione (reduced)** | N/A |
| 14 | **L-Carnitine HCl** | **L-Carnitine HCl** |
| 15 | **Linoleic Acid** | **Linoleic Acid** |
| 16 | **Linolenic Acid** | **Linolenic Acid** |
| 17 | **Progesterone** | **Progesterone** |
| 18 | **Putrescine 2HCl** | **Putrescine 2HCl** |
| 19 | **Sodium Selenite** | **Sodium Selenite** |
| 20 | **T3 (triiodo-L-thyronine)** | **T3 (triiodo-L-thyronine)** |

